# Deceiving The Big Eaters: *Salmonella* Typhimurium SopB subverts host cell Xenophagy through Akt-TFEB axis in macrophages

**DOI:** 10.1101/2022.02.03.479023

**Authors:** Ritika Chatterjee, Debalina Chaudhuri, Subba Rao Gangi Setty, Dipshikha Chakravortty

## Abstract

*Salmonella*, a stealthy facultative intracellular pathogen, harbors an array of host immune evasion strategies. This facilitates successful survival and replicative niches establishment in otherwise hostile host innate immune cells such as macrophages. *Salmonella* survives and utilizes macrophages for effective dissemination throughout the host causing systemic infection. One of the central host defense mechanisms in macrophages is bacterial xenophagy or macro-autophagy. Here we report for the first time that *Salmonella* pathogenicity island-1 (SPI-1) effector SopB is involved in subverting host autophagy through dual mechanisms. SopB is known to act as a phosphoinositide phosphatase and thereby can alter the phosphoinositide dynamics of the host cell. Here we demonstrate that this activity helps the bacterium escape autophagy by inhibiting terminal fusion of *Salmonella* containing vacuole (SCV) with both lysosomes and autophagosomes. We also report the second mechanism, wherein SopB downregulates overall lysosomal biogenesis through Akt- transcription factor EB (TFEB) axis. TFEB is a master regulator of lysosomal biogenesis and autophagy, and SopB restricts the nuclear localization of TFEB. This reduces the overall lysosome content inside host macrophages, further facilitating survival in macrophages and systemic dissemination of *Salmonella* in the host.

## Introduction

*Salmonella enterica* causes a range of infections, from self-limiting gastroenteritis to systemic typhoid fever in humans (Majowicz, Musto et al. 2010). *Salmonella* can gain access to non-phagocytic host cells through type 3 secretion system (T3SS) encoded by *Salmonella* pathogenicity island-1 (SPI-1) (Galan and Curtiss 1989, Collazo and Galan 1997) or directly phagocytosed by immune cells. Upon entering into the host cell, *Salmonella* resides in a unique membrane-bound compartment called *Salmonella*-containing vacuoles (SCVs)(Eswarappa, Negi et al. 2010, LaRock, Chaudhary et al. 2015). The stealthy pathogen injects several effector proteins by another T3SS encoded by SPI-2 to subvert the host innate defense pathways. Macrophages are the first line of host-innate immune cells that evade pathogen encounter. However, in the case of *Salmonella* Typhimurium pathogenesis, it is evident that the pathogen escapes the killing by macrophages and establish a replicative niche inside the cells to facilitate the systemic disease in susceptible hosts (Fields, Swanson et al. 1986, Leung and Finlay 1991, Das, Lahiri et al. 2009). Intracellularly, SCVs mature sequentially from the early (near periphery) to late SCVs (near juxtanuclear position) (LaRock, Chaudhary et al. 2015). In contrast, 10-20% of the SCV in epithelial cells rupture the vacuolar membrane and are either targeted to autophagic machinery or escape killing followed by hyper-proliferation (Perrin, Jiang et al. 2004, Malik-Kale, Winfree et al. 2012, Knodler, Nair et al. 2014). The cytosolic bacteria inside the macrophages encounter higher levels of stress such as high reactive oxygen species (ROS), reactive nitrogen species (RNS), antimicrobial peptides, metal starvation/toxicity, TLR/NLR signaling and a robust xenophagic machinery (Birmingham, Smith et al. 2006, Lahiri, Das et al. 2008, Das, Lahiri et al. 2009, Gogoi, Shreenivas et al. 2019). Therefore, bacteria must maintain the intact vacuolar membrane when residing inside the macrophages.

Several studies have shown that the *Salmonella* in SCV can escape autophagic machinery and proliferate inside hostile host cells when the vacuolar integrity is maintained. The autophagic adaptors, LC3B, SQSTM/p62, and NDP52 target cytosolic bacterium (Birmingham and Brumell 2006, Cemma, Kim et al. 2011), and Galectin-8 marks the damaged SCV membranes and then cleared through autophagy (Thurston, Wandel et al. 2012). Interestingly, the mechanisms are not clear yet how the intact SCV escapes the recruitment of autophagic machinery/adaptors and avoids fusion with the terminal lysosomes.

The fusion of autophagosomes with lysosomes majorly depends on the phosphoinositides regulation/conversion. The role of especially phosphatidylinositol 3-phosphate (PI(3)P) has been well characterized in autophagosome formation. In yeast, the dephosphorylation of PI(3)P by PI(3)P phosphatase, Ymr1, is an important event during the fusion of autophagosomes with the lysosomes (Cebollero, van der Vaart et al. 2012). The conversion of PI(3)P initiates the dissociation of the ATG machinery from the autophagosomal membranes leading to fusion with the terminal lysosomes. *Salmonella* also inhibits the fusion of SCVs with the lysosomes and autophagosomes when the vacuolar integrity is maintained. One of the effector molecules secreted by SPI-1, SopB, is known to act as phosphatidylinositol phosphatase, which converts PI(4,5)P to PI(3)P (Terebiznik, Vieira et al. 2002, Hernandez, Hueffer et al. 2004, Mason, Mallo et al. 2007, Mallo, Espina et al. 2008). Therefore, we hypothesized that SopB could be a potential candidate that inhibits the fusion of SCV with autophagosomes.

It is also well studied that SopB, even though a phosphatase, can activate Akt/protein kinase B by phosphorylating Ser473 residue and thus modulate downstream signaling in infected cells. SopB also inhibits apoptosis in the infected cells by activating Akt (Steele-Mortimer, Knodler et al. 2000, Knodler, Finlay et al. 2005, Raffatellu, Wilson et al. 2005). On the other hand, Akt is one of the critical modulators of Transcription Factor EB (TFEB) by phosphorylating TFEB at Ser467 residue. The phosphorylated TFEB (Ser467) shows reduced nuclear localization, resulting in downregulation of the genes under the TFEB promoter(Palmieri, Pal et al. 2017). TFEB positively regulates the set of genes under its promoter, termed as Coordinated Lysosomal Expression and Regulation (CLEAR) network in addition to autophagy genes(Settembre, Di Malta et al. 2011).

Interestingly, a previous study from our lab demonstrated that the number of lysosomes inside infected cells decreases upon *Salmonella* infection progression (Eswarappa, Negi et al. 2010). However, the mechanism as to how *Salmonella* depletes the number of lysosomes upon infection is unknown. Therefore, we hypothesized that SopB might play a dual role in subverting autophagy by (1) avoiding fusion with autophagosomes or lysosomes and (2) downregulating the overall lysosomes biogenesis by restricting transcription factor localization to the nucleus in the infected cells.

We show that *Salmonella* SopB employs dual mechanisms to modulate xenophagy. The first one is by accumulating PI(3)P on the SCV membranes, which successfully inhibits recruitment of autophagic adaptors on the SCV and the fusion of intact SCV with auto-phagolysosomes/lysosomes. The second mechanism employed by SopB is to restrict nuclear localization of TFEB, leading to downregulation of overall biogenesis of lysosomes and autophagosomes in macrophages. This gives the pathogen an advantage of survival as the ratio of SCV to lysosomes reduces. Thus, showing novel mechanisms that *Salmonella* Typhimurium SopB employs in subverting innate cellular defenses of the host that can serve as potential intervention targets.

## Results

### The intracellular proliferation of STM Δ*sopB* is attenuated in macrophages due to the recruitment of autophagic SNARE STX17

*Salmonella* SopB is one of the key effector proteins secreted by SPI-1 machinery into the host cell, and its role in inducing the uptake of the bacterium by epithelial cells has been well established (Raffatellu, Wilson et al. 2005). However, the role of SopB in the macrophages has not been fully elucidated. Therefore, we explored whether SopB has any role in phagocytosis and intracellular proliferation in macrophages. We found that in murine macrophage cell line RAW264.7, the STM Δ*sopB* mutant (knockout) had attenuated proliferation compared to *Salmonella* Typhimurium wildtype (STM WT) or STM Δ*sopB:sopB* (complemented strain). However, the percentage of phagocytosis remains unaltered compared to WT **(Figure 1A)**. The attenuated proliferation of STM Δ*sopB* mutant was also valid for human macrophage cell lines such as PMA stimulated U937 **(Figure 1B)** and Thp1 **(Figure 1D)** and mouse primary peritoneal macrophages **(Figure 1C)** as well. None of the strains exhibited any growth difference in *in-vitro* Luria Bertani (LB) broth (Figure S1A). Studies have shown that SopB is essential for the entry of *Salmonella* into the non-phagocytic host cells (Hume, Singh et al. 2017).

**Figure 1:**
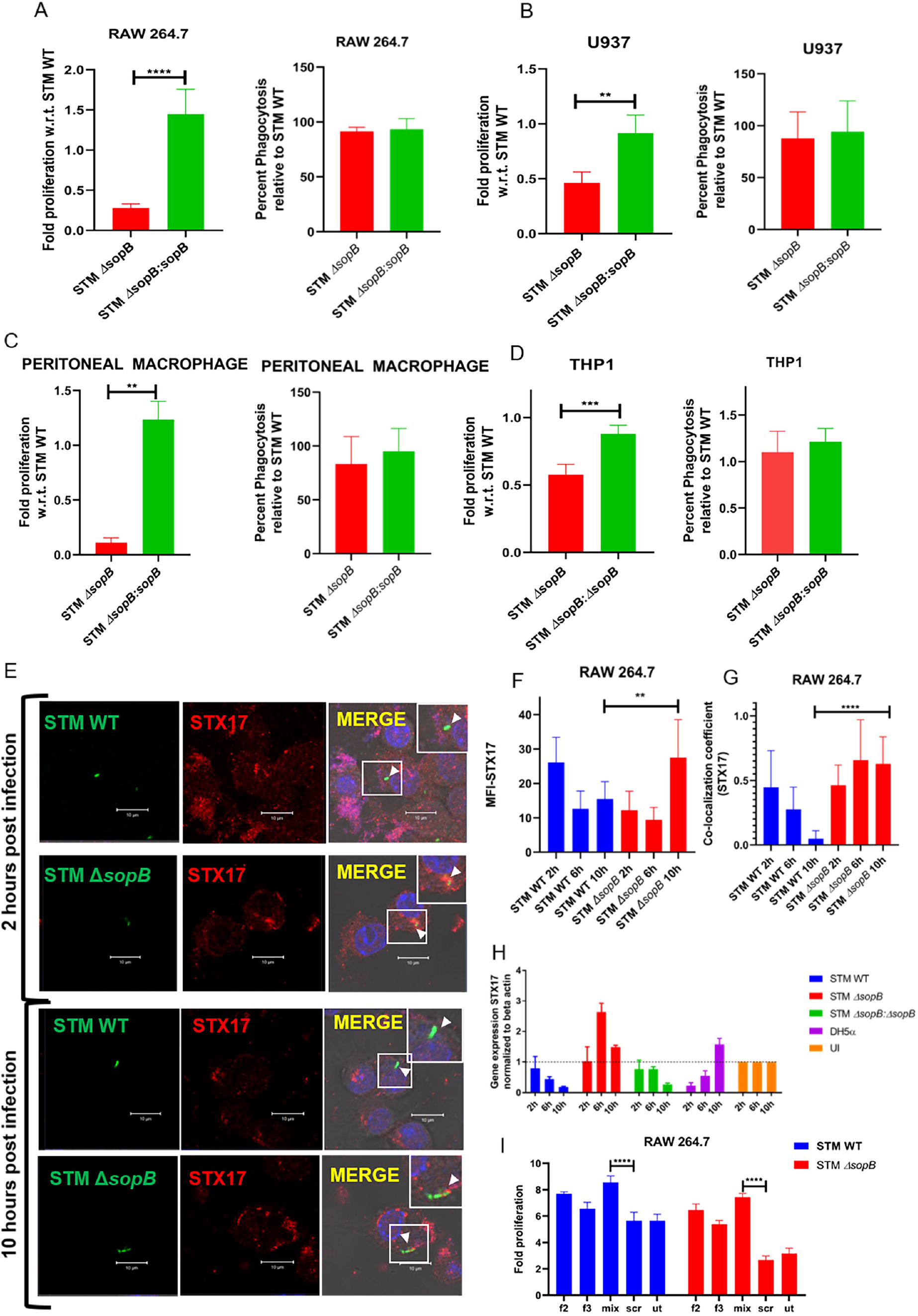
Intracellular proliferation of STM Δ*sopB* is attenuated in macrophages due to recruitment of autophagic SNARE STX17. Gentamicin protection assay performed in **(A)** murine macrophage RAW 264.7 cells with either STM WT or STM Δ*sopB* or STM Δ*sopB:sopB*, at 2h and 10h cells were lysed and plated to count CFU/mL and then fold proliferation is ratio of 16h/2h and, percentage invasion in RAW 264.7 cells was calculated 2h/pre-inoculum (p.i.)*100. Gentamicin protection assay performed in **(B)** human monocyte stimulated with PMA U937 cells **(C)** primary peritoneal macrophage cells **(D)** human monocyte stimulated with PMA Thp1. **(E)** Representative immunofluorescence images of infected RAW264.7 macrophages (MOI of 25) at 2h and 10hour in confocal laser scanning microscope (CLSM); Green-bacterial strain, Red-STX17, yellow -colocalization. Quantification of immunofluorescence images**(F)** Mean fluorescence intensity (MFI) and **(G)** colocalization coefficient was performed using ZEN 2.3 platform. **(H)** Quantitative RT-PCR to assess the levels of STX17 in infected RAW264.7 macrophages with STM WT or STM Δ*sopB* or STM Δ*sopB:sopB*. **(I)** Intracellular survival in STX17 knockdown cells. Scale bar in microscopic images is of 10μm and data is representative of one experiment with more than 50 cells analysed for each condition. All experiments were repeated at least three times N=3. Student’s unpaired t-test performed for statistical analysis, mean ± SEM/SD p<0.05, ** p<0.01, *** p<0.001.

Our data corroborated with Stévenin et al., that STM Δ*sopB* mutant maintains vacuolar integrity inside macrophages **(Figure S1N)** (Stévenin, Chang et al. 2019). This eliminates the possibility of encountering with high amount of reactive oxygen species, reactive nitrogen species and cationic antimicrobial peptides (Gogoi, Shreenivas et al. 2019). Therefore, we further analyzed the recruitment of autophagic markers onto the SCV membranes. One of the critical autophagic SNARE proteins is Syntaxin 17 (STX17), which mediates the fusion of autophagosomes with lysosomes (Nakamura and Yoshimori 2017). At 2 hours post-infection (2 hpi), we observed that the colocalization of STX17 onto the SCV membrane was comparable to STM WT and STM Δ*sopB* mutants. However, at later time points (10 hpi), there was a significant reduction of STX17 recruitment onto SCV membrane of STM WT and increased recruitment of STX17 onto SCV of STM Δ*sopB* mutants **(Figure 1E-G)**. This phenotype was rescued in complemented strain **(Figure S1B-E)**. We also have observed similar results in the case of human macrophages (U937) and mouse primary peritoneal macrophages upon infection with STM WT and STM Δ*sopB* mutants **(Figure S1G-K)**.

These observations indicate that SopB possibly inhibits the recruitment of autophagy machinery (STX17) onto the SCV membrane in macrophages. So, to further delve into the role of STX17, we carried out an intracellular fold proliferation assay of *Salmonella* in STX17 knockdown macrophages. We found that in the STX17 knockdown condition, the STM WT and STM Δ*sopB* mutants were able to survive significantly better than the scrambled control (**Figure 1I**). We also observed that the mean fluorescence intensity (MFI) of STX17 as visualized by confocal microscope was significantly lesser in the case of STM WT infection than STM Δ*sopB* mutant infection **(Figure 1F, S1E)**. Therefore, we analyzed the transcript levels of STX17 in infected RAW264.7 macrophages at different time points. We observed downregulation of STX17 transcript in the case of STM WT and STM Δ*sopB:sopB* (complemented strain) with the progression of infection but not in STM Δ*sopB* mutant infected macrophages. Interestingly, we did not observe any changes in the transcript levels of STX17 cognate SNAREs **(Figure S2A-C)**. These results indicated unknown underlying mechanisms might be employed by *Salmonella* SopB, which enables it to downregulate an autophagic SNARE at transcript levels.

### SopB inhibits the recruitment of other autophagic markers onto the SCV membrane

Next, we investigated whether the SopB plays a role in recruiting other autophagy adaptors such as LC3B and p62/SQSTM onto SCV, which are known autophagy adaptors, especially targeting intracellular pathogens(Wang, Yan et al. 2018). We found that similar to STX17, SopB also inhibits recruitment of both LC3B and p62/SQSTM onto SCV **(Figure 2A-B)** as observed by the colocalization coefficient **(Figure 2C-D)**. We also observed a significantly lesser number of puncta of p62/SQSTM per cell in STM WT infected macrophage cells than STM Δ*sopB* (Figure 2E), suggesting the overall induction of autophagy is significantly lesser in STM WT infected RAW264.7 macrophage cells. We also observed that recruitment of LC3B was pronounced in PFA treated STM WT (dead bacteria control) and in non-pathogenic *E. coli* DH5α **(Figure S2D)**. These observations strongly suggested that SopB is involved in restricting the induction of autophagy and inhibiting the recruitment of autophagy adaptor molecules on SCV in macrophages. We also observed a significant downregulation of adaptor proteins such as NDP52, LC3B, and p62/SQSTM in STM WT infected macrophages compared to STM Δ*sopB*. Therefore, we next sought to assess the protein levels of these adaptor molecules, and we found that most of them were upregulated in the case of STM Δ*sopB* mutants **(Figure 2I, S3A)**, suggesting a possible role of SopB in mediating the downregulation of autophagy adaptor proteins inside infected macrophages.

**Figure 2:**
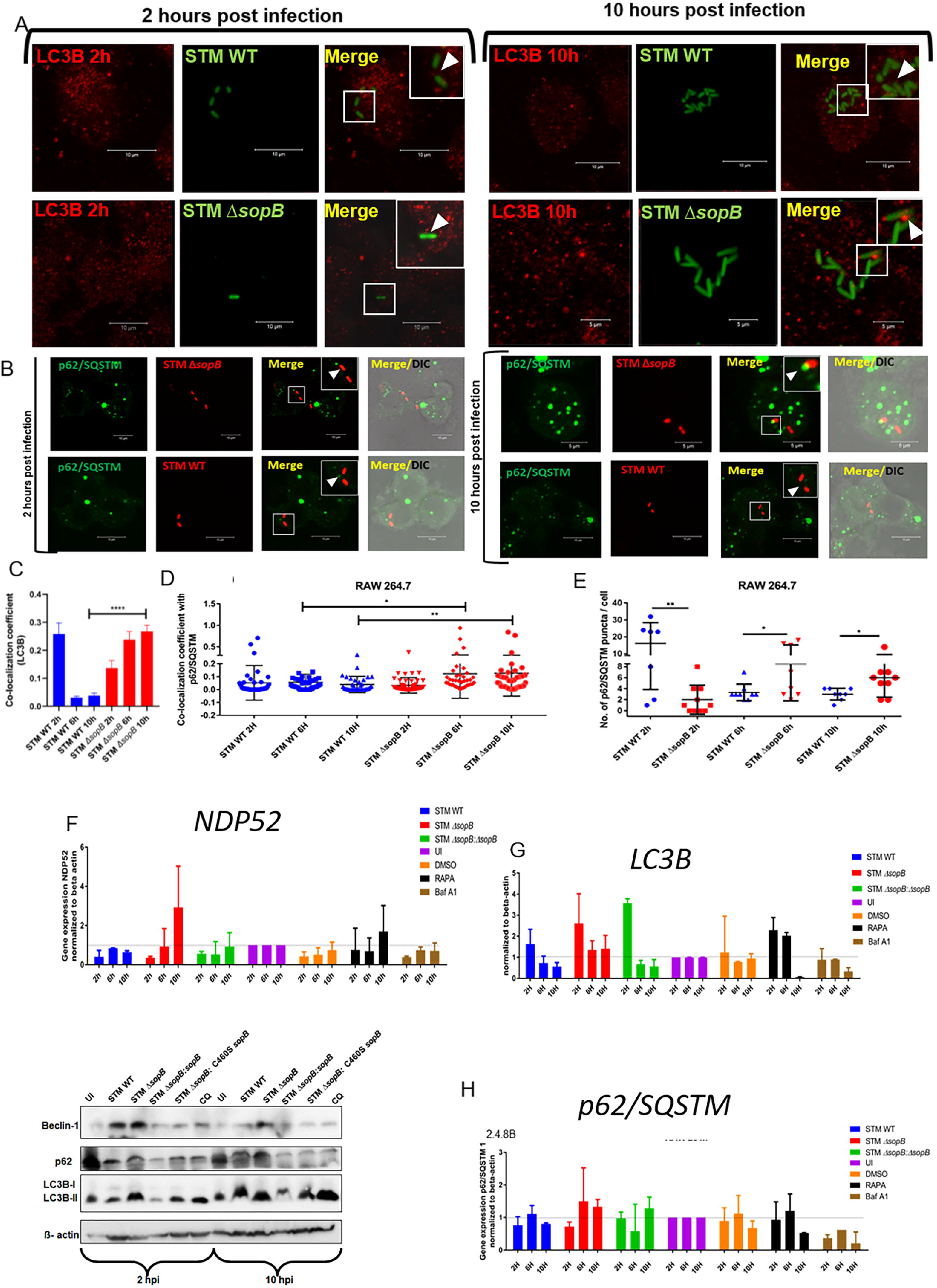
SopB inhibits recruitment of other autophagic markers onto the SCV membrane and also downregulate their levels at transcript and protein levels. Representative confocal microscopy images of RAW 264.7 cells infected with STM WT or STM Δ*sopB* and fixed at different time points, stained with **(A)** LC3B or **(B)** SQSTM/p62. **(C)** Quantification of LC3B colocalization at 2h and 10h. **(D)** Quantification of SQSTM/p62 colocalization coefficient and **(E)** puncta per cells were counted. Colocalization coefficient using Zen 2.3 platform. Quantitative RT-PCR of **(F)** NDP52, **(G)** LC3B and **(H)** p62/SQSTM, performed in the RAW264.7 murine macrophages infected with STM WT (blue) or STM Δ*sopB* (red) or STM Δ*sopB:sopB* (green), uninfected (purple), DMSO-vehicle control (orange), rapamycin treated (black) and bafilomycin A treated (brown). (I) Representative immunoblotting with Beclin1, p62 and LC3B. Scale bar in microscopic images is of 10μm and data is representative of one experiment with more than 50 cells analysed for each condition. All experiments were repeated at least three times N=3. Student’s unpaired t-test performed for statistical analysis, Mean ± SEM/SD * p<0.05, ** p<0.01, *** p<0.001.

### SopB inhibits fusion with autophagosomes and lysosomes by altering the phosphoinositide dynamics

SopB being a phosphoinositide phosphatase, is well known to alter the PI(3)P levels of SCV. Bakowski and colleagues had shown previously that the SopB is a crucial effector molecule that inhibits the fusion of SCV with lysosomes(Bakowski, Braun et al. 2010). Therefore, we hypothesized that similar mechanisms might help SCV avoid fusion with autophagosomes/auto-phago-lysosomes. We observed that the fusion events of SCV with autophagosome or lysosomes are reduced in STM WT infected macrophages compared to the STM Δ*sopB* mutants **(Figure 3A-B)**. This finding corroborated with the previous finding and provided newer insight that SopB plays a crucial role in dampening the fusion of SCV with autophagosomes.

**Figure 3:**
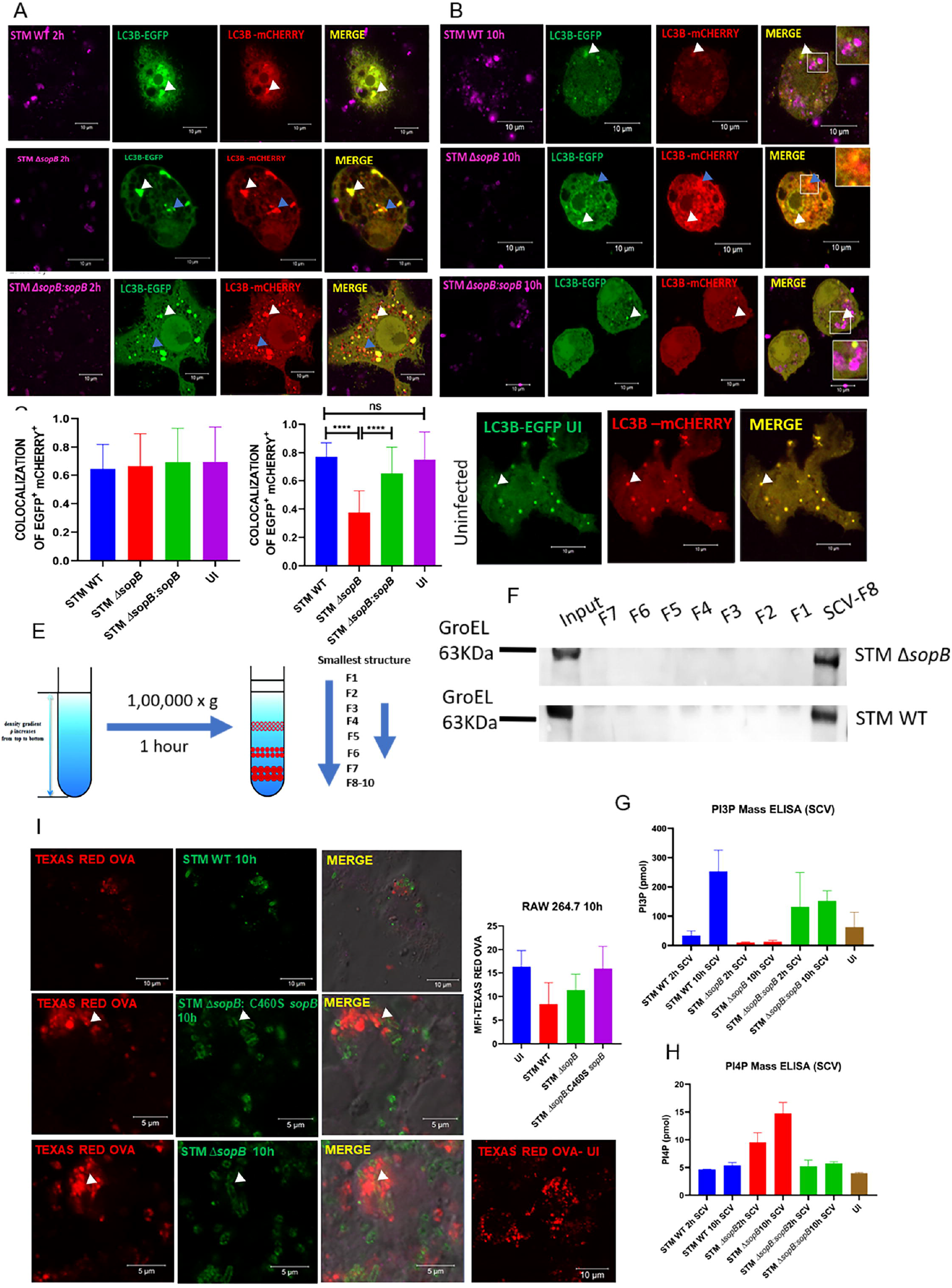
SopB inhibits fusion with autophagosomes and lysosomes by altering the phosphoinositide dynamics. Representative images of transiently transfected RAW264.7 macrophages with pDest EGFP mCherry LC3B tandem construct, further infected with STM WT or STM Δ*sopB* and cells fixed at **(A)** 2h and **(B)** 10h and further stained with anti-*Salmonella* to mark the bacterial localisation. Quantification of the colocalization of EGFP and mCherry at **(C)** 2h and **(D)** 10h using Zen 2.3 platform **(E)** Schematic of ultracentrifugation for isolation of *Salmonella* containing vacuoles (SCV)**; (F)** Immunoblotting with *Salmonella* GroEL with isolated fraction after ultracentrifugation to confirm that fraction 8-SCV fraction contains the purified bacterial population from infected cells. **(I)** To stain the terminal lysosomes 264.7 macrophages were treated with Texas red ovalbumin (OVA) for 45mins prior to infection with the STM WT or STM Δ*sopB or* STM Δ*sopB:* C460S *sopB*. **(J)** Quantification of the MFI using Zen 2.3 platform. **(G)** Representative PI(3)P and **(H)** PI(4)P mass ELISA plots of the isolated SCV fractions from ultracentrifugation. Scale bar in microscopic images is of 10μm and data is representative of one experiment with more than 50 cells analysed for each condition. All experiments were repeated at least three times N=3. Student’s unpaired t-test performed for statistical analysis, Mean ± SEM/SD p<0.05, ** p<0.01, *** p<0.001.

Interestingly, the overall fluorescence intensity and puncta of LC3B are also reduced in STM WT infected cells compared to STM Δ*sopB*. Next, we isolated the SCV from infected macrophages and performed mass ELISA, and we found significantly higher PI(3)P levels in STM WT isolated SCVs, which was completely abrogated in STM Δ*sopB* mutant SCVs. Concomitantly, the levels of PI4P onto the STM Δ*sopB* mutant SCV were significantly higher with the progression of infection **(Figure 3G-H)**. We also investigated the overall levels of PI(3)P and PI(4)P in the infected macrophages, and we observed that the levels of PI(3)P were overall higher in the case of STM WT infected cells than STM Δ*sopB* mutant. The levels of PI4P were only higher in cells infected with STM Δ*sopB* mutant at 10h post-infection **(Figure S3B-C)**. We further performed Texas red Ovalbumin (TROV) chase experiments with a catalytically dead SopB mutant (C460S) **(Figure S3E)** and the STM Δ*sopB* mutants to confirm our findings. In line with our previous observation, we find that SopB is a crucial effector inhibiting the fusion of SCV with autophagosomes and lysosomes **(Figure 3I-J)**. Together, all these data indicated that SopB is a key effector molecule from *S*. Typhimurium that helps inhibit the fusion of SCV with lysosomes and autophagosomes.

### SopB downregulates the overall lysosomal biogenesis by restricting the nuclear localization of TFEB into the nucleus

Since SopB is also a known Akt or Protein Kinase B modulator, it is involved in phosphorylation of Akt at Ser473 residue (Steele-Mortimer, Knodler et al. 2000, Knodler, Finlay et al. 2005, Raffatellu, Wilson et al. 2005). Interestingly, it is known that Akt can phosphorylate TFEB at Ser467 residue. The phosphorylated TFEB (Ser467) shows reduced nuclear localization, resulting in downregulation of the genes under the TFEB promoter (Palmieri, Pal et al. 2017). TFEB upregulates the set of genes under its promoter, termed as Coordinated Lysosomal Expression and Regulation (CLEAR) network and autophagy genes(Settembre, Di Malta et al. 2011). These genes are responsible for lysosomal biogenesis and autophagy-related processes under physiological conditions. We observed that upon infection with STM WT, there is an overall downregulation of the set of genes under TFEB **(Figure 4A-G)**. This was further confirmed in another human cell line-U937 as well. We have also confirmed the same with confocal microscopy, where we observed that the TFEB nuclear localization was reduced in the case of STM WT as compared to catalytically dead mutant or STM Δ*sopB* mutants where we observed an increased colocalization of TFEB into the nucleus similar to a non-pathogenic *E. coli* DH5α **(Figure 4H-K)**.

**Figure 4:**
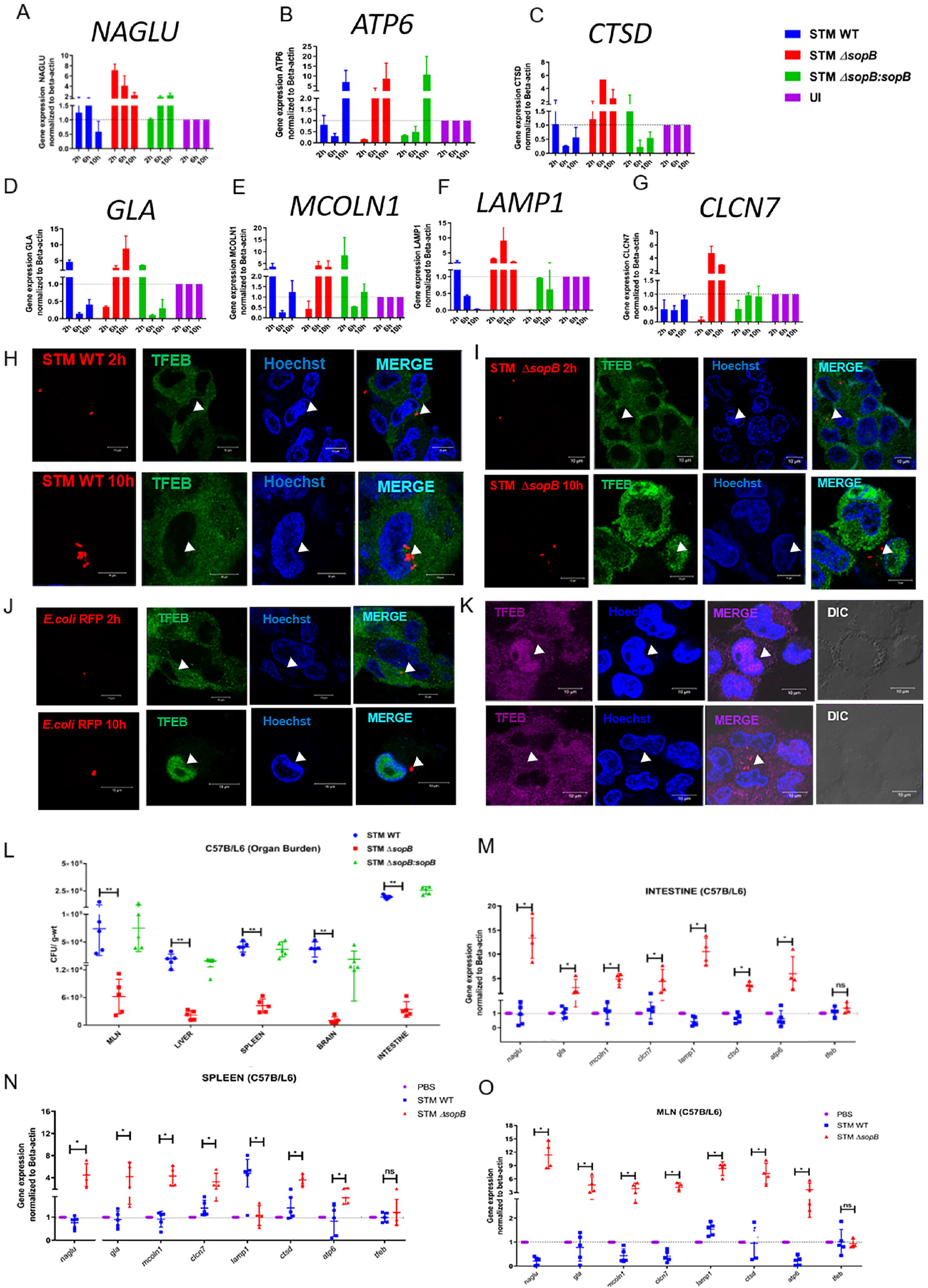
SopB downregulates the overall lysosomal biogenesis by restricting the nuclear localisation of TFEB into the nucleus. Representative quantitative RT-PCR for lysosomal biogenesis gene **(A)** NAGLU, **(B)**ATP6, **(C)** CTSD, **(D)** GLA, **(E)** MCOLN1 **(F)** LAMP1 **(G)** CLCN7 from RAW264.7 macrophages infected with STM WT or STM Δ*sopB* or STM Δ*sopB:sopB*. Representative confocal images of infected RAW264.7 macrophages with mentioned bacteria in red panel and stained with anti-TFEB (Green) antibody to assess its localisation inside the cells. To mark the nucleus, we have used Hoechst staining RAW264.7 murine macrophages infected with **(H)** STM WT **(I)** STM Δ*sopB* **and (N)** STM Δ*sopB:sopB* RAW264.7 murine macrophages, 10h post infection with infection with a catalytically dead SopB harbouring STM. Scale bar in microscopic images is of 10μm and data is representative of one experiment with more than 50 cells analysed for each condition. All experiments were repeated at least three times N=3. Student’s unpaired t-test performed for statistical analysis, Mean ± SEM/SD * p<0.05, ** p<0.01, *** p<0.001. C57BL/6 mice of 4-6 weeks were gavaged with 10^7^ CFU/mL of bacteria and 3 days post infection, mice were sacrificed **(L)** organ bacterial burden was calculated and the organs tissue RNA isolated for quantitative RT-PCR in **(M)** Intestine **(N)** Spleen **(O)** MLN. All quantitative RT-PCR data are representative of one biological replicate, and mean ± SD * p<0.01, ** p<0.001, *** p<0.0001.

We were then interested in deciphering if the same phenomena occur in the animal model system; we observed that the overall bacterial burden in organs was less in STM Δ*sopB* mutants than WT or complemented strain. So, we further assess the levels of the genes under TFEB promoter in different tissues colonized by *Salmonella* of the infected mice. We observed that SopB is involved in the overall downregulation of these genes at a tissue level, reducing the overall lysosomal biogenesis and autophagy flux in tissue-specific levels **(Figure 4L-O)**. Together, these results suggest that SopB inhibits overall lysosomal biogenesis and autophagic pathways through the TFEB-Akt axis. Our study reveals dual mechanisms employed by SopB to subvert host-mediated xenophagy in macrophages.

## Discussion

Facultative and obligated intracellular pathogens are known to modulate the host endocytic pathways to establish their unique replicative niche in host cells. Several bacterial molecules regulate host endocytic or defense pathways (Weber and Faris 2018). Xenophagy of the bacterium is one of the crucial pathways employed by host cells to keep the bacteria infection in check (Wileman 2013). *Coxiella burnetii* utilizes autophagosomes that provide the bacteria with a source of the membrane, facilitating its survival within host cells (Romano, Gutierrez et al. 2007, Vazquez and Colombo 2010). However, in the case of *Legionella pneumophila*, which subverts autophagy and cross-talk between ER-mitochondria by cleaving the syntaxin 17 with effector molecule Lpg1137 (Arasaki, Mikami et al. 2017). Therefore, establishing a successful niche inside the host cell is pathogen-specific.

*Salmonella* proliferates within-host innate immune cells like macrophages and utilizes the niche to establish systemic infection (Fields, Swanson et al. 1986, Leung and Finlay 1991, Das, Lahiri et al. 2009). Even though studies have shown more significant participation of SPI-2 effectors in intracellular survival and proliferation, well-orchestrated cross-talk between SPI-1 and SPI-2 effectors cannot be undermined (Lou, Zhang et al. 2019). Several diverse canonical roles of SopB (SPI-1 effector) are well dissected in epithelial cells, where studies show that SopB is associated with facilitating hyper-replication once the bacterium is in the cytosol of epithelial cells. However, there is a dearth in understanding the role of SopB in macrophages. Macrophages are professional phagocytes that can take up bacteria without induction and mounts more robust xenophagy than epithelial cells (Germic, Frangez et al. 2019), indicating that *Salmonella* might subvert or utilize host xenophagy depending on the cell type it is residing.

We report a model (Figure 4) for the first time that, SopB subverts xenophagy in host macrophages through a dual mechanism. Firstly, it alters the PI(3)P levels of the SCV membrane to inhibit its fusion with auto-phagolysosomes and lysosomes. Intracellular pathogens often find their haven inside the modified phagosome or vacuolar environment of host cells to remain hidden from innate defense pathways. Whence there is a possibility that due to altered PI(3)P levels, SCV remains as a hidden organelle inside the host cells. Thus, autophagy machinery fails to identify it, resulting in escaping an otherwise robust innate defense mechanism.

Our lab and others have shown that intracellular pathogens modulate the lysosomal biogenesis in host cells(Sachdeva and Sundaramurthy 2020). We here report a second mechanism employed by SopB to downregulate the overall lysosomal and autophagosomal biogenesis through the TFEB-Akt axis, reducing overall lysosomal content in infected macrophages, thereby giving an upper hand to an intracellular pathogen because of the number of active lysosomal to SCV ratio reduces. Our study also draws attention towards therapeutic strategies that can be further delved into to assess the ability of small molecule inhibitors (against SopB) or activators of TFEB, which might help reduce systemic *Salmonella* infection through macrophages.

## Material and Methods

### Bacterial strains and growth condition

*Salmonella enterica* serovars Typhimurium (STM) wild type strain ATCC SL13344, STM *ΔsopB*, STM *ΔsopB*: *sopB* (expressing SopB through a pWSK29-low copy number plasmid) were a kind gift from Prof. Michael Hensel, Abteilung Mikrobiologie, Universität Osnabrück, Osnabrück, Germany. *E. coli* DH5α were cultured in Luria broth (LB-Hi-media) with constant shaking (175rpm) at 37°C. Ampicillin or kanamycin was used wherever required. The bacteria were tagged with mCherry with pPFPV 25.1 plasmids for immunofluorescence studies. Site-directed mutagenesis was done by Phusion polymerase (New England Bio Labs) using a primer as previously described(Liebl, Qi et al. 2017) to generate STM *ΔsopB*: C460S *sopB* (expressing C460S SopB through a pWSK29-low copy number plasmid).

### Cell culture protocol

The cells RAW264.7 murine macrophages were cultured in DMEM - Dulbecco’s Modified Eagle Medium (Sigma) supplemented with 10% FBS (Gibco) at 37°C in a humidified incubator (Panasonic) with 5% CO2. Prior to each experiment, the cells were seeded onto the required plate either with a coverslip (for confocal fluorescence microscopy) or without (for intracellular survival assay, qRT-PCR and western blotting) at a confluency of 80-90%.

Human monocytes/macrophages U937 and Thp1 were cultured in RPMI-Roswell Park Memorial Institute media supplemented with 10% FBS (Gibco) at 37°C in a humidified incubator (Panasonic) with 5% CO2. Prior to each experiment, the cells were seeded with Phorbol-12-myristate-13-acetate (PMA-Sigma) at a concentration of 20ng/mL onto the required plate either with a coverslip (for confocal fluorescence microscopy) or without (for intracellular survival assay, qRT-PCR and western blotting) at a confluency of 80-90%.

Peritoneal macrophages isolation was performed as previously described (Zhang, Goncalves et al. 2008). Briefly, 4-6 weeks C57BL/6 mice were injected with Brewer’s thioglycollate media (Hi-Media) in the peritoneal cavity, and after 4-5 days post-intra-peritoneal (i.p.) injection, the cells were harvested from the peritoneal cavity. Cells were then seeded, and 24hour post-harvesting experiments were performed.

### Gentamicin protection assay

The cells were then infected with *Salmonella* Typhimurium (STM) (strain SL1344), STM *ΔsopB*, STM *ΔsopB*: *sopB*, STM WT PFA fixed and DH5α. at MOI of 25 for confocal experiment. Upon infecting the RAW264.7 cell-line, the plate was centrifuged at 700-900 rpm for 5 mins to facilitate the adhesion and then incubated for 20mins at 37°C and 5% CO_2_. Post-incubation, the bacteria containing media were removed, and wells were twice washed with PBS, and fresh media was added containing 100μg/mL gentamicin, incubated for 1 hour at 37°C and 5% CO_2_. Following this, the media was removed, washed with PBS twice and 25μg/mL gentamicin-containing media was added and incubated for different time points at 37°C and 5% CO_2_. Time points selected for confocal microscopy, qRT-PCR and immunoblotting were 2 hours, 6 hours and 10 hours post-infection. The time points were 2 hours and 10 hours for intracellular survival assay.

### Confocal Microscopy

After appropriate hours of incubation post-infection with STM-WT, STM *ΔsopB*, STM *ΔsopB*: *sopB*, STM WT PFA fixed and DH5α. The cells on coverslips were washed thrice with PBS and fixed with 3.5 % paraformaldehyde for 10-15mins. Then cells were washed twice with PBS and incubated with a specific antibody (α-STX 17 (Protein-Tech) or α-LC3B(Novus Biologicals) in a blocking buffer containing 2 % BSA and 0.01% saponin for 3 hours at room temperature (RT) or overnight at 4°C. Following this, the cells were washed twice with PBS and incubated with an appropriate secondary antibody tagged with fluorochrome for 1 hour at RT. The coverslips were then mounted onto a clean glass slide with mounting media and antifade agent; after the mounting media dried, it was sealed with clear nail polish and imaged under a confocal scanning laser microscope (Zeiss 880 microscope, at 63X oil immersion, 2x-3x zoom).

### Bacterial enumeration for intracellular survival assay

After appropriate hours of incubation post-infection with STM-WT, the mammalian cells were lysed by 0.1 % Triton-X 100. Then the lysate was plated onto *Salmonella*-Shigella (SS) Agar plate at appropriate dilutions. Percentage invasion and fold proliferation were then calculated with the following formula.

Percent invasion = CFU at 2h / CFU of Pre-Inoculum * 100

Fold Proliferation = CFU at 10h/ CFU at 2h.

### RNA isolation and quantitative RT PCR

RNA isolation was performed from transfected cells after appropriate hours of infection with STM WT at MOI of 10 or from tissue samples mesenteric lymph nodes (MLN), spleen and intestine (infected C57BL/6) by using TRIzol (Takara) reagent according to manufacturers’ protocol. Quantification of the RNA was done in NanoDrop (Thermo-Fisher Scientific). To check for RNA quality, the isolated RNA was also run on 2% agarose gel, and 3μg of RNA has subjected to DNase 1 treatment at 37°C. The reaction was then stopped with the addition of EDTA, and heated at 65°C for 10mins. As per the manufacturer’s protocol, the cDNA was synthesized by a cDNA synthesis kit (Takara). Quantitative real-time PCR was done using SYBR/ TB green (TAKARA) RT-PCR kit in BioRad qRT-PCR system. The reaction was set up in a 384 well plate with three replicates for each sample. The expression levels of the gene of interest were measured using specific RT primers (Table S1). Gene expression levels were normalized to beta-actin as an internal control.

### Transient Transfection

RAW 264.7 cells were seeded at a 50-60% confluency 12 hours prior to transfecting using either PEI (1:2 -DNA: PEI) or Lipofectamine 3000 (Thermo-fisher) as per manufacturer’s protocol. Approximately 300-500ng of plasmid DNA/well (ratio 260/280 ∼1.8-1.9) was used for transfection in 24well plate, and 1-2μg of plasmid DNA/well was used for 6well plates. List of plasmids used is given in Table S2. Cells were then incubated for 8hours at 37°C in a humidified incubator with 5% CO2; after that, the media containing transfecting DNA and reagents were removed, and cells were further incubated for 48 hours in complete media DMEM +10% FBS. Cells were then either harvested for further analysis or infected with the required MOI.

### Isolation of *Salmonella* containing vacuole (SCV) by Ultracentrifugation

The isolation of SCV was performed as previously described (Luhrmann and Haas 2000). Roughly 50 million RAW264.7 cells infected with *S*. Typhimurium SL1344 strain were used for subcellular fractionation of SCVs. At 2 hr, 6 hr and 10 hr p.i., cells were washed thrice with ice-cold PBS and scrapped into a 15 ml centrifuge tube using a rubber cell scraper. The cells were centrifuged at 1000 rpm for 7 min, and the cell pellets were suspended in ice-cold homogenization buffer (250 mM sucrose, 20 mM HEPES (pH 7.2), 0.5 mM EGTA and protease inhibitor cocktail (Roche) and transferred to a Dounce Homogenizer with a tight-fitting pestle on ice to break the cells. Approximately 30 strokes were applied until almost 90% of the cells were broken without breaking the nuclei. The intact cells and nuclei were pelleted at 400 x g for 3 min. The resulting supernatant was collected in a fresh tube to yield the post-nuclear supernatant (PNS). The PNS was brought to a final concentration of 39% sucrose and layered on to 2 ml 55% sucrose, which was layered onto a 65% sucrose cushion in a 13.2 ml open-top Beckman ultracentrifuge tube followed by the addition of 2 ml 32.5% and 2 ml 10% sucrose solutions. All sucrose solutions (w/v) were prepared in 20 mM HEPES (pH 7.2) and 0.5 mM EGTA. The PNS layered on sucrose gradient was then subjected for ultracentrifugation in a swinging bucket rotor for 1 hr at 100000 x g at 4°C. The fractions of 1 ml each were collected from top to bottom. Pooled fractions 8-10 were adjusted very slowly to a final sucrose concentration of 11% with homogenization buffer without sucrose and layered on a 15% Ficoll cushion (5% sucrose, 0.5 mM EGTA and 20 mM HEPES pH 7.2). The samples in an open-top Beckman ultracentrifuge tube were spun at 18000 x g for 30 minutes in a Beckman SW 41 Ti rotor at 4°C. The supernatant was discarded, and the pellet was resuspended in an 11 ml homogenization buffer. The samples were spun again at 18000 x g for 20 min in a Beckman SW 41 Ti rotor at 4°C, and the resulting pellet was labelled as an “SCV” fraction. The pelleted SCV fractions were resuspended in 200μL of homogenization buffer.

### PI(3)P and PI(4)P Mass ELISA

Isolated SCV fraction from infected RAW264.7 macrophages were further processed for lipid isolation and the isolated lipids were quantified using PI(3)P and PI(4)P mass ELISA kits (Echelon Biosciences) as per manufacturer’s protocol.

### Texas Red Ovalbumin Pulse chase experiment

RAW 264.7 cells were seeded at a 50-60% confluency 12 hours prior to treatment with Texas red Ovalbumin (Thermo-Fischer Scientific) at a concentration of 50μg/mL for 30minutes at 37°C in a humidified incubator with 5% CO2. Next, media was removed and fresh medium with stationary phase bacteria (10-12hours old) at a MOI of 25 was added to the cells and further incubated for 25mins in humidified incubator. At indicated timepoints cells were washed twice with 1X PBS and fixed with 3.5% paraformaldehyde for 15minutes. Cells were images under microscope after staining with anti-*Salmonella* antibody (Zeiss LSM 880) using 63X oil immersion objective lens and images were analysed with ZEN Black 2009 software by Zeiss.

### *In-vivo* experiments

6 weeks old C57BL/6 mice were infected by oral gavaging of 10^7^ CFU of STM WT, STM *ΔsopB, or* STM *ΔsopB: sopB*. 5 days post-infection, mice were sacrificed, and organs such as the liver, spleen, MLN, brain and intestine were plated onto SS agar, and tissue samples from spleen MLN and intestine were also used for RNA isolation and further analysis.

### Immunoblotting

After appropriate hours of infection with STM WT at MOI of 10, the media was removed, and the cells were washed twice with PBS. Cells were then harvested using a sterile scraper and centrifuged at 1500 rpm for 10 mins, 4°C. Cell lysis was done by RIPA buffer for 30mins on ice, followed by estimation of total protein using Bradford protein estimation method. Polyacrylamide Gel Electrophoresis (PAGE) was done by loading 35μg of protein from whole cell lysate, then transferring onto 0.45μm PVDF membrane (GE Healthcare). The membrane was blocked using 5% skimmed milk (Hi-Media) in TTBS for 1h at RT and then probed with specific primary and secondary HRP conjugated antibodies. The membrane was developed using ECL (Bio-rad), and images were captured using ChemiDoc GE healthcare. All densitometric analysis was performed using the Image J Platform.

### Statistical analysis

Each experiment has been independently performed at least 3 times (as mentioned in figure legends). Confocal data sets were analysed and quantified in Zen 2.3 platform by Zeiss. The data sets were analysed by unpaired student’s t-test by GraphPad Prism 8.4.3 software, and *p-values* are indicated in the figures and legends for reference. The results are either expressed as mean ± SEM or mean ± SD as indicated in the legends. Data obtained from *in-vivo* mouse experiments were analysed by Mann-Whitney *U* test from GraphPad Prism 8.4.3 software

## Acknowledgment

Divisional and Departmental Confocal Facility, Departmental Real-Time PCR Facility, and Central Animal Facility at IISc are duly acknowledged. Mr. Punith, Mrs. Saima and Ms. Navya are acknowledged for their help in image acquisition. Ms. Sukeerthi, Ms. Nishi and Ms. Rhea are acknowledged for their technical help.

## Availability of data and materials

All data generated and analysed during this study, including the supplementary information files, have been incorporated in this article. The data is available from the corresponding author on request.

## Declarations

### Ethics statement

All the animal experiments were approved by the Institutional Animal Ethics Committee, and the Guidelines provided by National Animal Care were strictly followed. (Registration No: 48/1999/CPCSEA)

