## SUPPLEMENTAL for "Deceiving The Big Eaters: *Salmonella* Typhimurium SopB subverts host cell Xenophagy through Akt-TFEB axis in macrophages"

SUPPLEMENTARY:

Figure S1

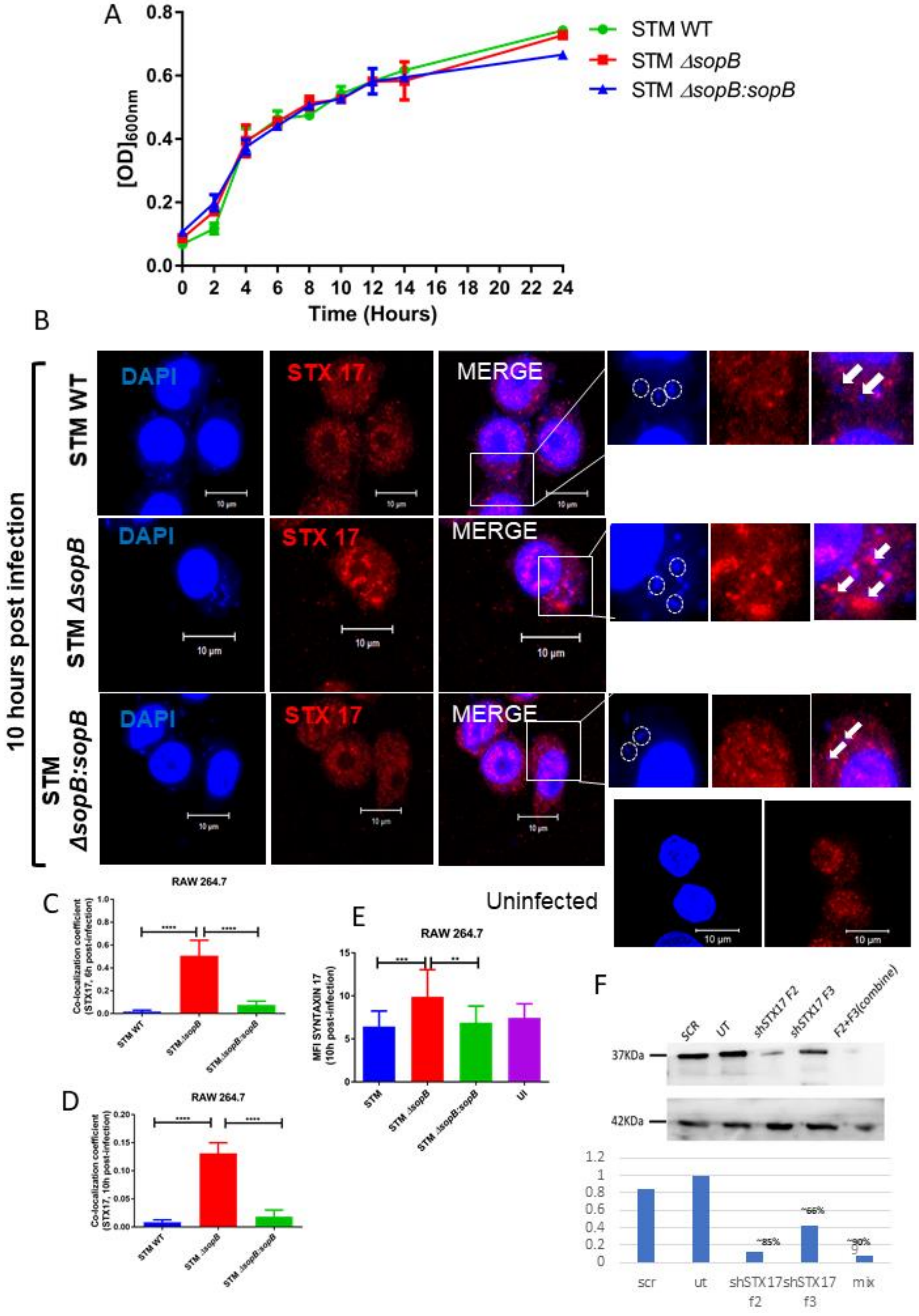

Figure S1

G

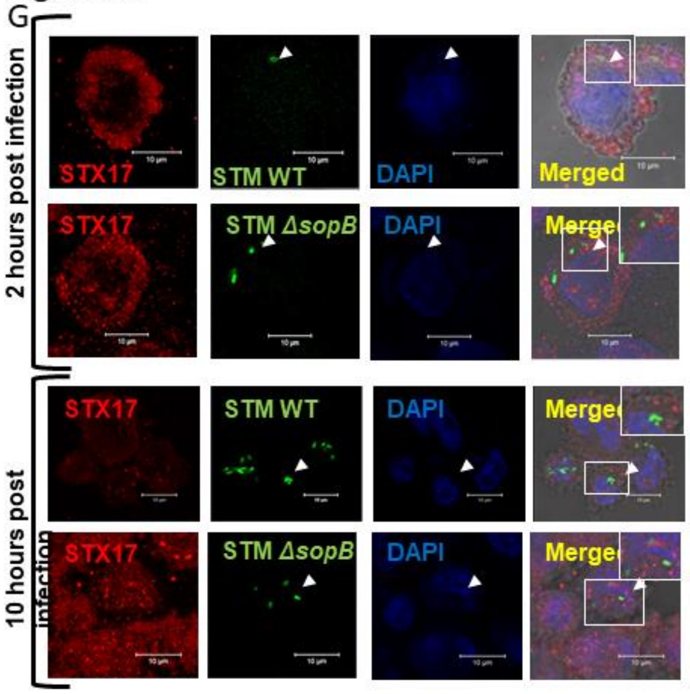

H

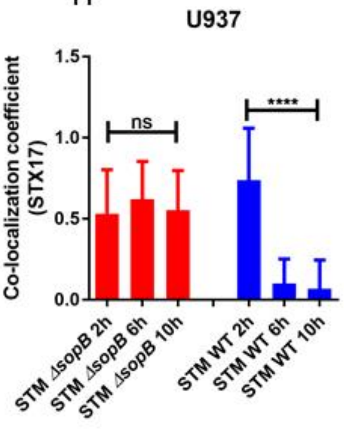

I STM WT 10h

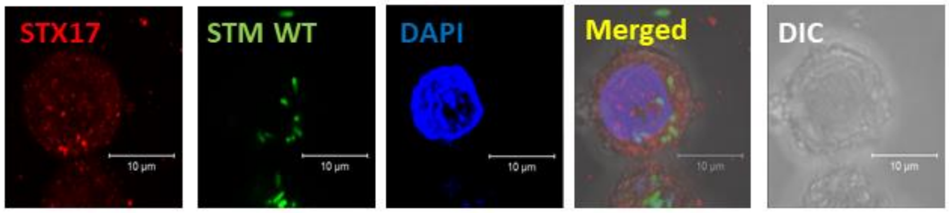

J

STM *ΔsopB* 10H

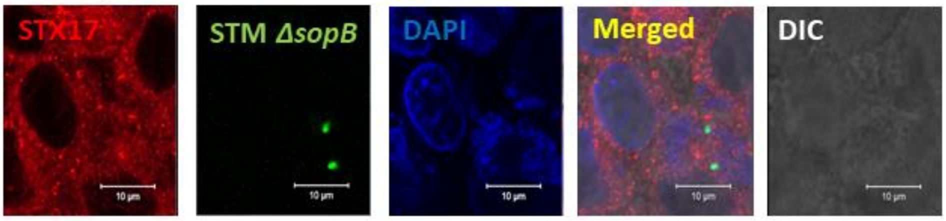

Figure S1

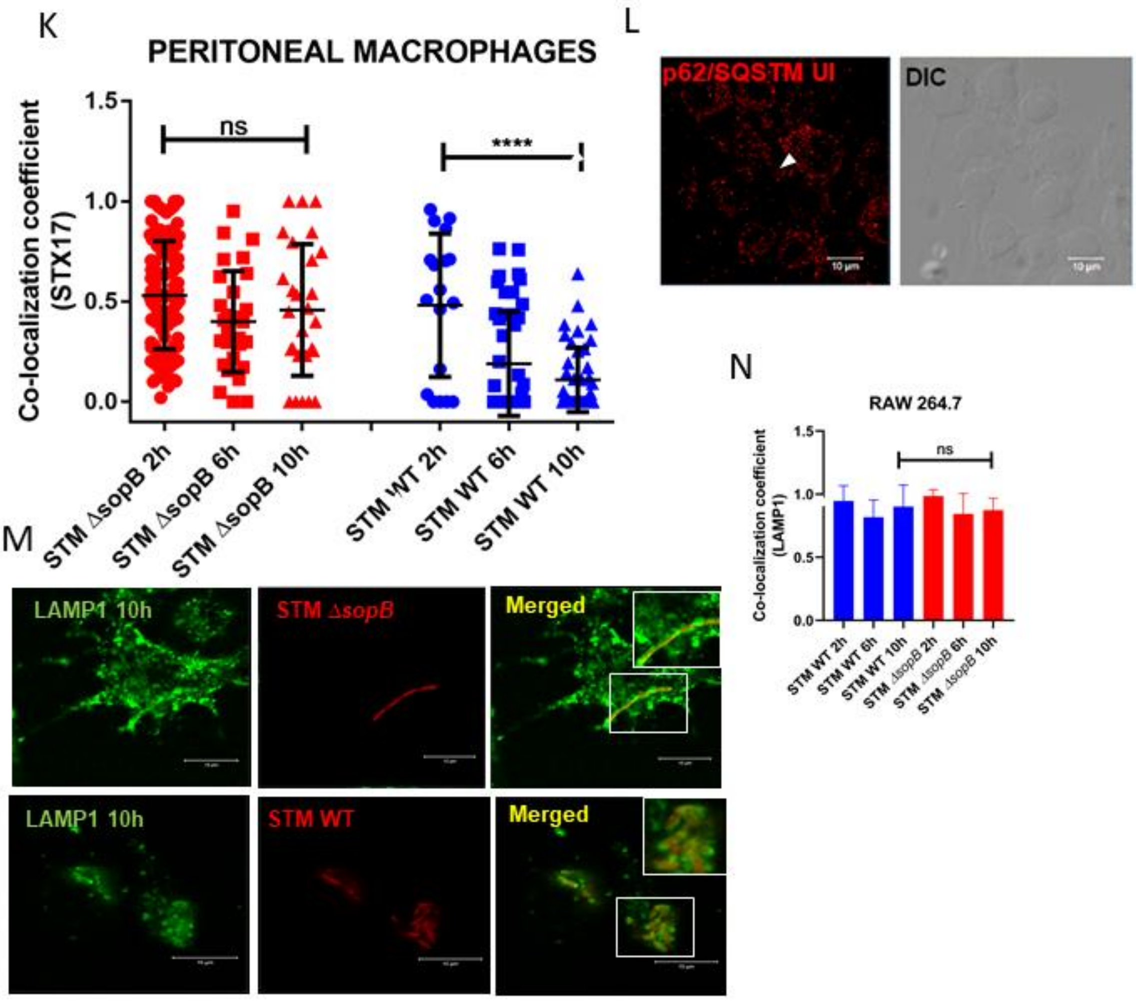

**Figure S1:** (A) Growth curve analysis of all three strains in LB media.

(B) Representative images of infected RAW264.7 macrophages, stained with anti-STX17 at 10h post infection, the bacteria marked with DAPI.

Quantification of colocalization co-efficient using ZEN2.3 platform at (C) 6h post infection and (D) 10h post infection. (E) Quantification of mean fluorescence intensity using ZEN 2.3 platform; (F) Immunoblotting of STX17- knockdown cells and densitometric analysis of immunoblot.

(G) Representative images of infected human monocyte/macrophages U937, stained with anti-STX17 at 10h post infection, the bacteria tagged with mCherry. (H) Quantification of colocalization co-efficient using ZEN2.3 platform at different time-points of infection.

Representative images of infected with (I) STM WT and (J) STM Δ*sopB* primary macrophages isolated from peritoneal cavity of C57BL/6 mice, stained with anti-STX17 at 10h post infection, the bacteria tagged with mCherry. (K) Quantification of colocalization co-efficient using ZEN2.3 platform at different time-points of infection.

(L) uninfected cells stained with anti- p62/SQSTM.

(M) Representative images of infected RAW 264.7 murine macrophages, stained with anti-LAMP1 at 10h post infection, the bacteria tagged with mCherry. (N) Quantification of colocalization co-efficient using ZEN2.3 platform at different time-points of infection.

Figure S2

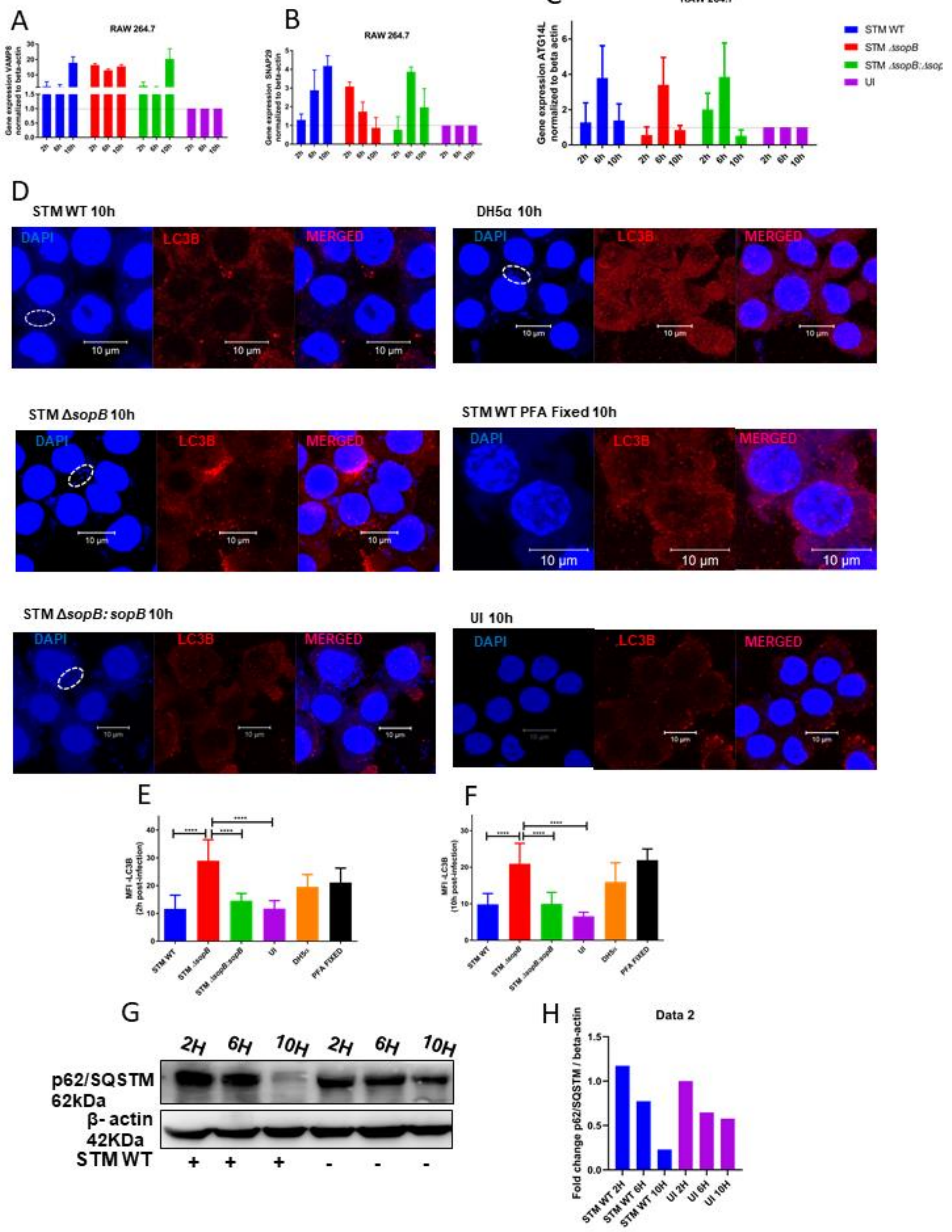
A
RAW 264.7
Gene expression VAMP8 normalized to beta-actin
B
RAW 264.7
Gene expression SNAP29 normalized to beta-actin
C
RAW 264.7
Gene expression ATG14L normalized to beta actin
STM WT
STM ΔsopB
UI
D
STM WT 10h
DH5α 10h
STM ΔsopB 10h
STM WT PFA Fixed 10h
STM ΔsopB: sopB 10h
UI 10h
DAPI
LC3B
MERGED
10 μm
E
MFI -LC3B (2h post-infection)
F
MFI -LC3B (10h post-infection)
STM WT
STM ΔsopB
STM ΔsopB:sopB
UI
DH5α
PFA FIXED
G
2H
6H
10H
p62/SQSTM 62kDa
β- actin 42KDa
STM WT
H
Data 2
Fold change p62/SQSTM / beta-actin
STM WT 2H
STM WT 6H
STM WT 10H
UI 2H
UI 6H
UI 10H

Figure S2: Quantitative RT-PCR to assess the levels of STX17 cognate SNAREs (A) VAMP8, (B) SNAP29 and (C) ATG14L in infected RAW264.7 macrophages with STM WT or STM Δ*sopB* or STM Δ*sopB:sopB*.

(D) Representative images of infected RAW 264.7 murine macrophages, stained with anti-LC3B at 10h post infection with mentioned bacteria. Quantification of mean fluorescence intensity of LC3B using ZEN 2.3 platform at (E) 2h post infection and (F) 10h post-infection.

(G) Immunoblotting with p62/SQSTM with different timepoint of infection in murine macrophages RAW264.7, (H) Densitometric quantification of the immunoblot with ImageJ.

Figure S3

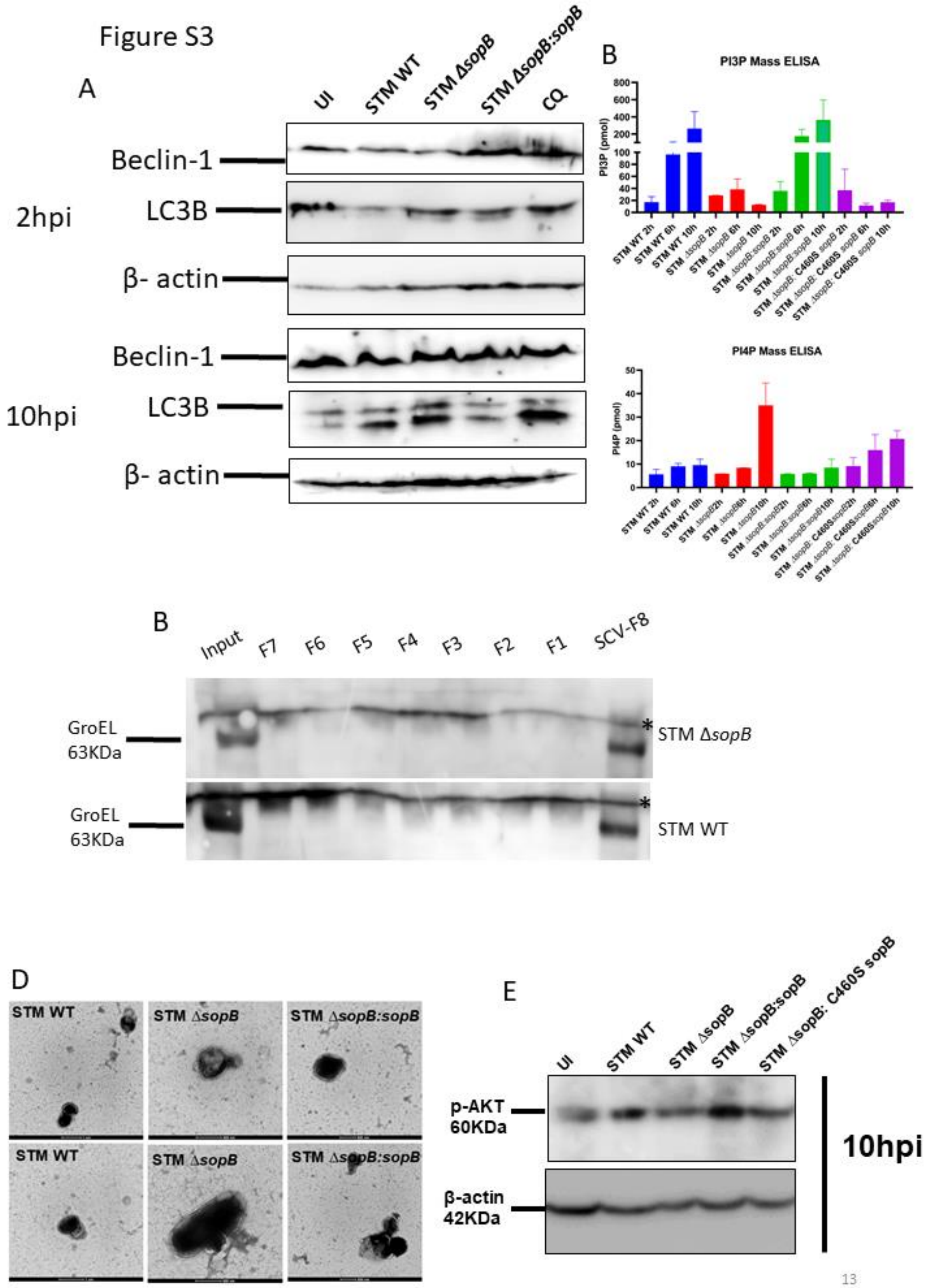
A
UI
STM WT
STM ΔsopB
STM ΔsopB:sopB
CQ
Beclin-1
LC3B
β- actin
2hpi
10hpi
B
PI3P Mass ELISA
PI3P (pmol)
PI4P Mass ELISA
PI4P (pmol)
B
Input
F7
F6
F5
F4
F3
F2
F1
SCV-F8
GroEL
63KDa
STM ΔsopB
STM WT
D
STM WT
STM ΔsopB
STM ΔsopB:sopB
E
UI
STM WT
STM ΔsopB
STM ΔsopB:sopB
STM ΔsopB: C460S sopB
p-AKT
60KDa
β-actin
42KDa
10hpi

Figure S3: (A) Immunoblotting of beclin 1 and LC3B at different timepoint of infection in murine macrophages RAW264.7.

(B) Mass ELISA of PI(3)P and PI(4)P of the infected murine macrophages RAW264.7 cells with STM WT, STM Δ*sopB,* STM Δ*sopB:sopB* and STM Δ*sopB:* C460S *sopB.*

(C) Full length immunoblotting of GroEL for isolated SCV fractions. * -Indicate non-specific bands

(D) Representative electron micrograph images of the isolated SCV with ultracentrifugation.

(E) phospho-Akt immunoblot with infected RAW 264.7 macrophages at 10 hours post infection with indicated strains.

### Figure S4

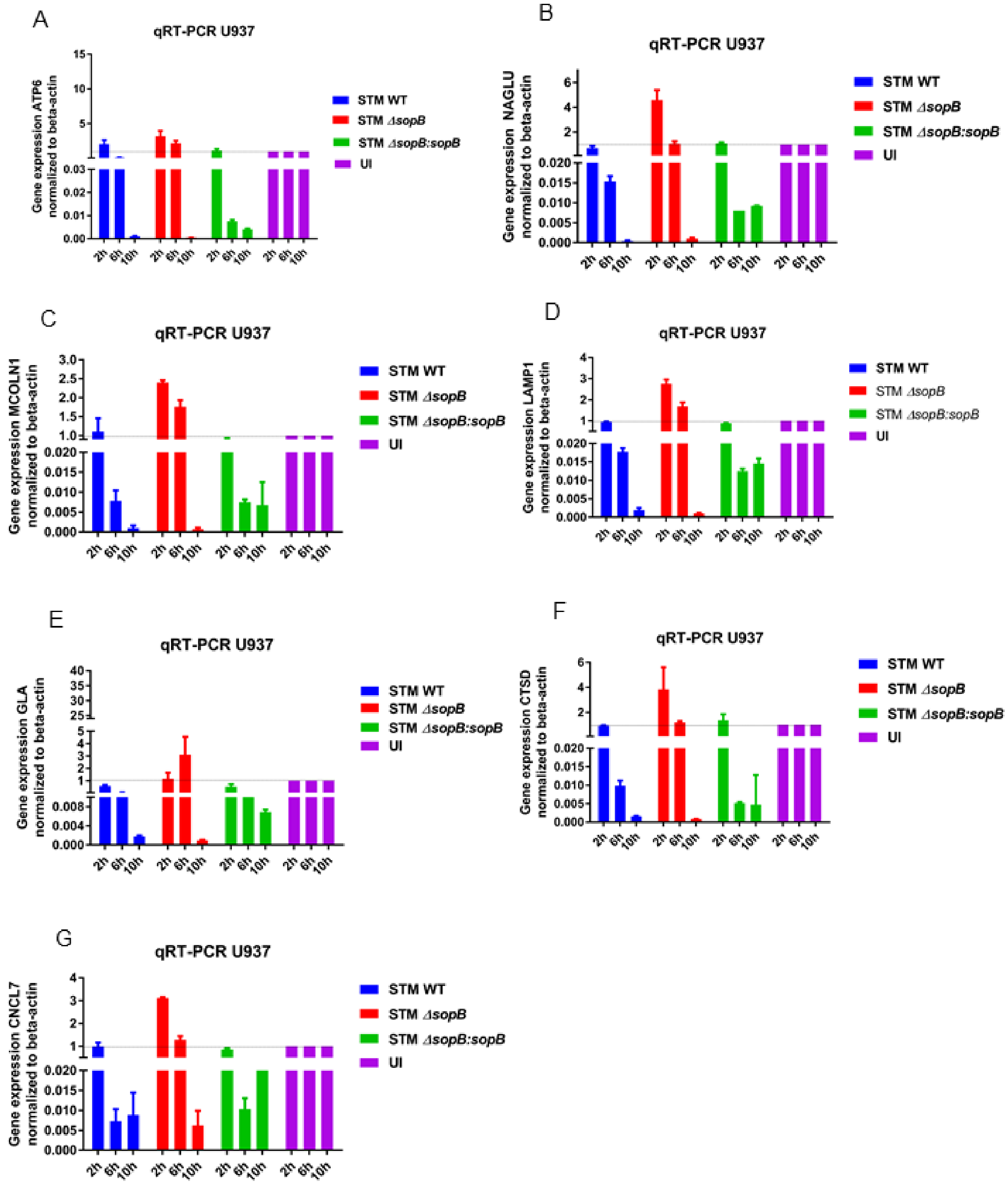
A
qRT-PCR U937
Gene expression ATP6 normalized to beta-actin
B
qRT-PCR U937
Gene expression NAGLU normalized to beta-actin
C
qRT-PCR U937
Gene expression MCOLN1 normalized to beta-actin
D
qRT-PCR U937
Gene expression LAMP1 normalized to beta-actin
E
qRT-PCR U937
Gene expression GLA normalized to beta-actin
F
qRT-PCR U937
Gene expression CTSD normalized to beta-actin
G
qRT-PCR U937
Gene expression CNCL7 normalized to beta-actin
STM WT
STM ΔsopB
STM ΔsopB:sopB
UI
2h
6h
10h

Figure S4: Representative quantitative RT-PCR for lysosomal biogenesis gene **(A)** ATP6**,** **(B)**NAGLU**,** **(C)** MCOLN1**,** **(D)** LAMP1, **(E)** GLA, **(F)** CTSD **(G)** CLCN7 from human monocyte/macrophages U937 infected with STM WT or STM Δ*sopB* or STM Δ*sopB:sopB* normalised to uninfected.

| **Table S1: List of primers (5'- 3')** | |
|---|---|
| *LC3B FP* | GGGGCAGGCGGGTGATTAT |
| *LC3B RP* | ACTTCGGAGATGGGAGTGGA |
| *STX 17 FP* | CTAGGCGGGAGGTGTTTCTGTT |
| *STX 17 RP* | TCAAGCCTGCGTAACTTCACC |
| *ATG 14 FP* | AGACTTCTGGGAGAGACCCTG |
| *ATG 14 RP* | AGTCCCCGTTGTTTGGTAGG |
| *SNAP 29 FP* | AACCTAGATGAGCTGTCCGT |
| *SNAP 29 RP* | AGAGTTCCTCCTCCGTGTCT |
| *P62/SQSTM FP* | GGGACAGCCAGAGGAACAGAT |
| *P62/SQSTM RP* | TCCTTCCTGTGAGGGGTCTA |
| *NDP52 FP* | CTGGCTGCTGTTCCTGGACT |
| *NDP52 RP* | GATGCCAATCCAGTCCTTGC |
| *β -ACTIN FP* | CAGCAAGCAGGAGTACGATG |
| *β -ACTIN RP* | GCAGCTCAGTAACAGTCCG |
| *hu-ACTB-FP* | CATGTACGTTGCTATCCAGGC |
| *hu-ACTB-RP* | CTCCTTAATGTCACGCACGAT |
| *hu-LAMP1 FP* | TTCAAGGTGGAAGGTGGC |
| *hu-LAMP1 RP* | CCAGAGAAAGGAACAGAGGC |

| | |
|---|---|
| *hu-CTSD FP* | ACCTCGTTTGACATCCACTATG |
| *hu- CTSD RP* | AGGATGCCATCGAACTTGG |
| *hu-GLA FP* | CGCTTCATGTGCAACCTTG |
| *hu- GLA RP* | TCTGCCTTCTGAATCTCTTTGG |
| *hu-NAGLU FP* | CTTTCAATGAGATGCAGCCAC |
| *hu- NAGLU RP* | TCAGCAAACAGGTCCAGAAC |
| *hu- MCOLN1 FP* | CCACATCCAGGAGTGTAAGC |
| *hu- MCOLN1 RP* | CCAGCCATTGACAAATTCCAG |
| *hu-ATP6 FP* | AGCTATTGACACTTCCTGGTG |
| *hu- ATP6 RP* | AGCAAACTGAACAGGTCACC |
| *HU- CLCN7 FP* | GCTCTGTGATTGTGGCTTTC |
| *HU- CLCN7 RP* | TTCAGTGACGTTGACCTTCC |
| *Hgprt_ for_mouse* | GCAGCGTTTCTGAGCCATTG |
| *Hgprt_rev_mouse* | TCATCGCTAATCACGACGCT |
| *Clcn7_for_mouse* | CTGGGGTATGGTTTGATGCC |
| *Clcn7_rev_mouse* | TAATGGGGTTCTCTCAGGCT |
| *Atp6_for_mouse* | AGTCCGGCTTACAGCTAACA |
| *Atp6_rev_mouse* | GGAGGGTGAATACGTAGGCT |
| *Gla_for_mouse* | ACGATCTGCGACAAATCAGC |
| *Gla_rev_mouse* | CGCAGGATATGTTCCTTGGC |
| *Ctsd_for_mouse* | CTATAAGCCGGCGACCTCTG |
| *Ctsd_rev_mouse* | TGCAATGAATGGAGGGGACC |
| *Tfeb_for_mouse* | AGCGAGAGCTAACAGATGCT |
| *Tfeb_rev_mouse* | GGATGGTGCCTTTGTTCCAG |

| | |
|---|---|
| *Naglu_for_mouse* | CTGTGACTTCCACGTGTGTG |
| *Naglu_rev_mouse* | TGCCTCTACCTTCTGACTGC |
| *Lamp1_for_mouse* | TCAACCTCTGGACACTTGGC |
| *Lamp1_rev_mouse* | GCATTACGTGAGCTGATGCC |
| *Mcoln1_for_mouse* | GACCAAGACCCACATCCAGG |
| *Mcoln1_rev_mouse* | CCGTTCCCAGAGGCTGATTT |

Table S2 Plasmids used:

| **S. No.** | **Plasmids** | **Source** |
|---|---|---|
| 1 | pDest-EGFP-mCherry-LC3B- tandem construct | Prof. S. Vijaya |
| 2 | shRNA – Syntaxin 17 TRCN0000379933 (F2) | MISSION shRNA -pLKO.1 SIGMA |
| 3 | shRNA – Syntaxin 17 TRCN0000379634 (F3) | MISSION shRNA -pLKO.1 SIGMA |
| 4 | shScramble - non target control | MISSION shRNA -pLKO.1 SIGMA |
